# A molecular inventory of the faecal microbiomes of 23 marsupial species

**DOI:** 10.1101/2025.08.31.672981

**Authors:** Kate L. Bowerman, Rochelle M. Soo, Pierre-Alain Chaumeil, Michaela D. J. Blyton, Mette Sørensen, Disan Gunbilig, Maika Malig, Moutusee Islam, Julian Zaugg, David L. A. Wood, Ivan Liachko, Benjamin Auch, Mark Morrison, Lutz Krause, Birger Lindberg Møller, Elizabeth H. J. Neilson, Philip Hugenholtz

## Abstract

Despite the recent expansion of culture-independent analyses of animal faecal microbiomes, many lineages remain understudied. Marsupials represent one such group, where despite their iconic status, direct sequencing-based analyses remain limited. Here we present a metagenomic and metabolomic exploration of the faecal microbiomes of 23 *Diprotodontia* marsupials, producing a reference set of 3,868 prokaryotic and 12,142 viral metagenome-assembled genomes, the majority (>80%) of which represent novel species. As with other animals, host phylogeny is the primary driver of microbiome composition, including distinct profiles for two eucalyptus folivore specialists (koalas and southern greater gliders), suggesting independent solutions to this challenging diet. Expansion of several bacterial and viral lineages were observed in these and other marsupial hosts that may provide adaptive benefits. Antimicrobial resistance genes were significantly more prevalent in captive than wild animals likely reflecting human interaction. This molecular dataset contributes to our ongoing understanding of animal faecal microbiomes.

**Impact statement:** Despite their ecological and evolutionary importance, marsupials remain underrepresented in microbiome research. Here, we present the most extensive faecal microbiome dataset to date for this group, encompassing metagenomic, metabolomic, and proximity ligation data from 23 marsupial species. As in other animals, we find the microbial community structure reflects the host species, and some marsupials carry expanded sets of certain microbial lineages indicative of within-host evolution. This work substantially expands the genomic landscape of host-associated microbes and viruses in a poorly studied mammalian clade.

**Data summary:** Raw read data, prokaryotic MAGs ≥50% complete with ≤10% contamination are available via the European Nucleotide Archive under project PRJEB89408. The full set of viral genomes, clustered protein database and metabolite data (raw and processed) are available via https://doi.org/10.48610/14e37e9. Prokaryotic MAGs are also available via https://figshare.com/s/87443d80817f57aadc16.

## Introduction

Marsupials are celebrated Australian fauna, however, their microbiomes remain underexplored. Marsupials comprise seven taxonomic orders, of which the *Diprotodontia* harbour the majority of iconic species including koalas (*Phascolarctus cinereus*), wombats (e.g. *Vombatus ursinus*) and kangaroos (e.g. *Osphranter rufus*) ^1^. However, only 27 of the 151 extant *Diprotodontia* species ^2^ have had their faecal microbiomes characterised using culture-independent methods, and only 14 of these have been analysed using metagenomics (**Table S1**) ^3–29^. To our knowledge, no other omic techniques have been applied to marsupial microbiomes and no culture-independent analyses of marsupial viral communities have been reported.

Here, we applied metagenomics and metabolomics to faecal samples from 23 *Diprotodontia* species, including two eucalyptus folivore specialists, koalas and southern greater gliders, to contribute to our knowledge of microbial species associated with marsupials. Eucalyptus leaves are an unusual diet due to their low nutrient availability and plethora of complex and often toxic secondary metabolites that deter herbivores, including terpenes, tannins, cyanogenic glycosides, flavonoids and formylated phloroglucinol compounds ^30–32^. Furthermore, there is a high degree of biochemical variation between leaves of the estimated 150-200 *Eucalyptus* species that marsupials feed on and koalas and greater gliders have different feeding preferences ^31,33–35^. Here, we find this reflected in their faecal metagenomes and metabolomes indicating independent solutions to mastering this challenging diet. Despite the evolutionary divergence and geographical isolation of marsupials from other mammals over most of their history ^36^, prokaryotic taxonomic novelty of the analysed faecal microbiomes was only observed at the genus and species level, suggesting that readily discoverable higher level microbial diversity in the animal gut is reaching saturation.

## Results and discussion

### Comparison of marsupial faecal microbiomes

To extend our knowledge of the marsupial gastrointestinal microbiome, we characterised 94 faecal samples from 23 host animal species (and one subspecies *Trichosurus vulpecula* subsp. *fuliginosus*, golden brushtail possum) via metagenomic and metabolomic analyses (**Tables S2 & S3**). These species represent 17 genera and 6 families in the order *Diprotodontia*, of which this study provides the first metagenomic data for 13 species, 9 genera and 1 family (**Fig. 1**). A further set of 23 faecal samples from nine of these species was investigated using proximity-ligation sequencing to provide links between microbial hosts and their viruses and plasmids ^37^ (**Table S3**).

**Figure 1.**
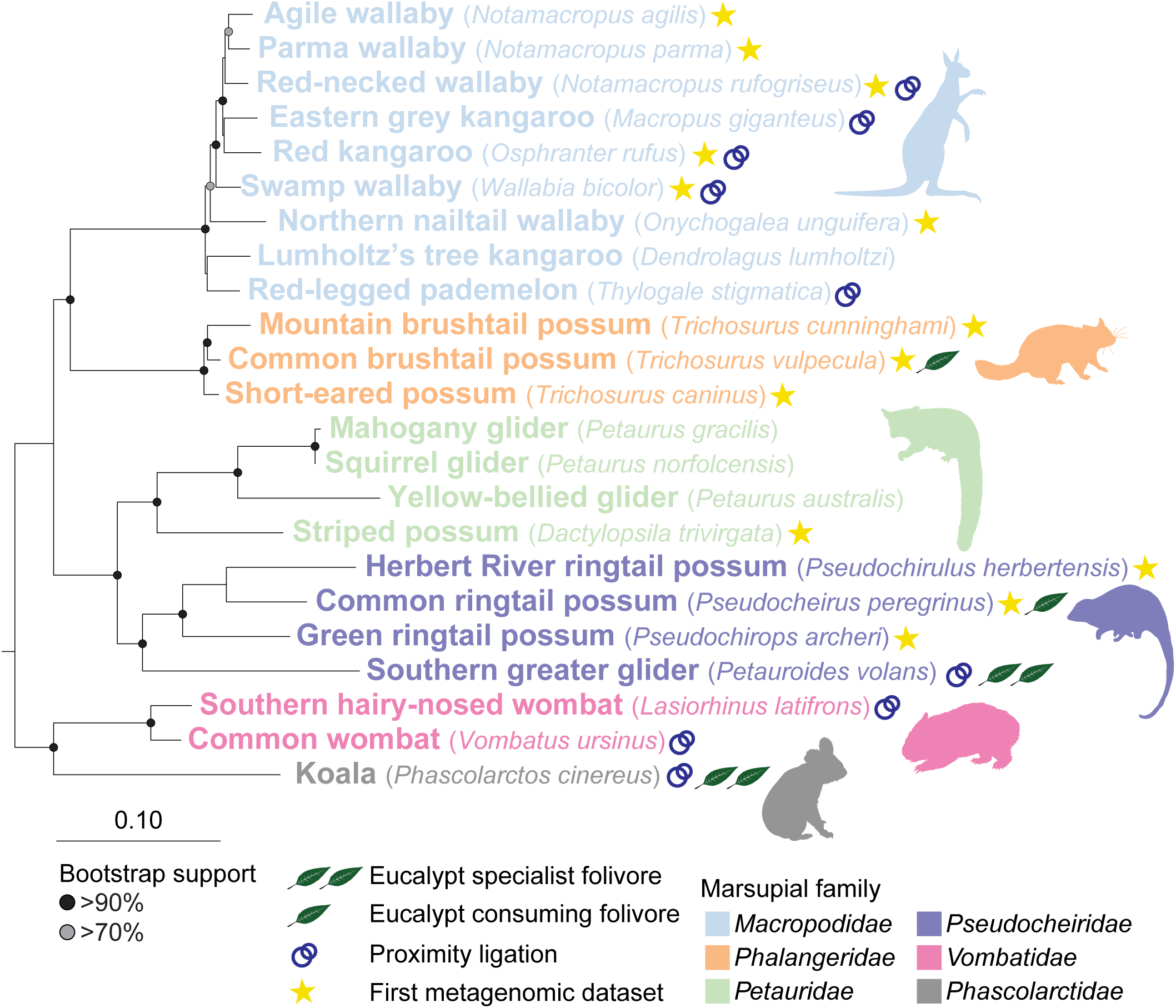
A maximum-likelihood phylogenetic tree of the 23 marsupial species used in this study (subspecies *Trichosurus vulpecula fuligonosus* was excluded due to the closeness to *Trichosurus vulpecula*) and an outgroup (*Dromiciops gliroides*) was generated from a concatenated alignment of nine genes (four chromosomal, three mitochondrial and two ribosomal) ^107^. Stars indicate hosts for which this study provides the first metagenomic data. Interlocked circles denote samples sequenced using both standard metagenomic and proximity ligation sequencing.

We compared faecal microbiomes based on their taxonomic (prokaryotic and viral), functional (KEGG and CAZy) and metabolomic profiles (**Fig. 2**). In all cases, host phylogeny (marsupial family) was the primary contributor to variance ranging from 12% (viral taxa) to 31% (CAZymes). This is consistent with previous reports of host phylogeny contributing the most variance to faecal profiles in animals, indicative of phylosymbiosis between animals and their microbiomes ^38–40^.

**Figure 2.**
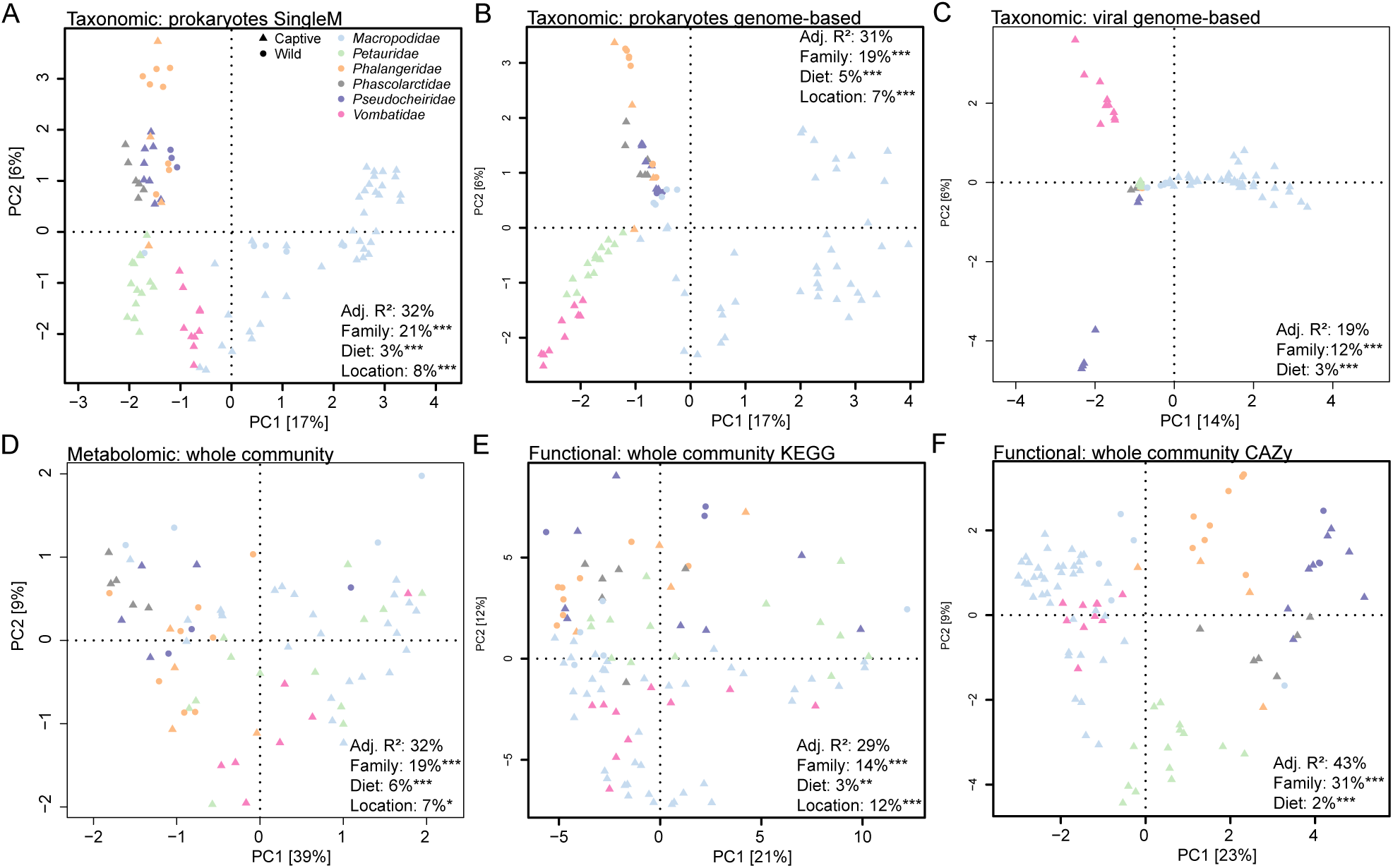
Principal component analysis based on **a,** marker gene-based prokaryotic community **b,** genome-based prokaryotic community **c,** genome-based viral community **d,** metabolomic **e,** KEGG-based functional and **f,** CAZy-based functional profiles. Taxonomic relative abundances were centred log-ratio transformed and functional RPKM values were log_2_ transformed prior to PCA. Metabolomic data as described in the methods. Samples are coloured based on their marsupial host family. Results of redundancy analysis incorporating host family, diet (grazer, browser, omnivore, fungivore), gut type (hindgut, foregut), location and captivity status displayed per dataset.

Sample location (wildlife sanctuary or region where samples were obtained) and diet (grass-dominant grazers, foliage-dominant browsers, omnivores, fungivores) also contributed significantly to microbiome variance (**Fig. 2**), consistent with other mammals ^38,41^. Wild/captive status was not significantly correlated with faecal profiles, despite known differences in some marsupial species, such as higher incidence of the bacterial family *Muribaculaceae* in captive *vs* wild koalas ^10^. This may reflect sampling bias as fewer scat samples were obtained from wild animals in our study (19% wild *vs* 81% captive). Macropod microbiomes were taxonomically distinct from those of other marsupial species forming a primary divide using principal component analysis (**Fig. 2A-C**), driven by differences in the classes *Clostridia* and *Bacteroidia*, or their associated phages (**Fig. S1A-C, Tables S4-S6**). This may be in part due to macropods being the sole foregut fermenting family in the dataset. However, gut morphology (foregut *vs* hindgut) did not explain additional variance in the faecal profiles beyond that captured by the host family (**Fig. 2**) and the distinction was less apparent in functional and metabolomic PCA plots (**Fig. 2D-F, Tables S7-S9**). The functional profiles of the grazing families *Macropodidae* and *Vombatidae* overlapped, particularly evident in the CAZyme profiles where they were separated from other families along the primary component axis (**Fig. 2F**). Multiple xylan and mannan-degrading CAZymes were amongst the top contributors to this separation, likely used for digestion of hemicelluloses that are more abundant in grasses compared to leaves and other browse (**Fig. S1F, Table S9**). *Macropodidae* and *Vombatidae* also harboured the highest faecal microbial diversity, with browsing animals significantly lower (**Fig. S2**), as seen previously in African herbivores ^42^.

We then compared the faecal profiles of specialist (koala, southern greater glider) and generalist eucalyptus feeders (common and golden brushtail and common ringtail possums) to look for signs of shared adaptations to this challenging diet. Despite similarities in the context of all marsupials investigated (**Fig. 2**), analysis in isolation revealed taxonomic, functional and metabolomic differences between these groups (**Fig. S3, Tables S10-S17**). Multiple eucalypt-derived terpenes positively associated with the specialist feeders (**Fig. S3B, Table S11**) indicative of the high rates of eucalypt ingestion in these hosts. However, the profiles of the two specialist hosts were distinct. For example, koala faeces were higher in essential oil metabolites and greater glider faeces in wax-derived compounds (**Fig. S3E, Table S15**), likely reflecting feeding preferences. Koalas favour leaves of the subgenus *Symphyomyrtus* that are often rich in complex oils and greater gliders prefer the leaves of the subgenus *Monocalyptus* that have thick wax layers ^43–45^.

### Prokaryotic communities of the marsupial gut

We individually assembled and binned the 94 standard Illumina shotgun metagenomes (averaging ∼3.2 Gbp per sample) obtaining a total of 908 medium to high quality (≥70% complete, <5% contaminated, N50 >10 kb) prokaryotic metagenome-assembled genomes (MAGs) (**Table S4**). An additional 2,960 medium to high quality prokaryotic MAGs were obtained from the 23 proximity ligation metagenomes (averaging ∼56 Gbp per sample). The combined set of 3,868 MAGs (3,828 bacterial and 40 archaeal) represent 2,412 species, 659 genera, 134 families, 59 orders, 27 classes and 19 phyla, of which 1,951 species (80%) and 58 genera (9%) are new to GTDB (release 09-RS220) (**Table S18**) ^46^. The lack of higher taxonomic rank novelty (family and above) is notable especially as this study represents the first metagenomic data for many marsupial species (**Table S1**), suggesting that microbial diversity recoverable from animal faeces as MAGs is beginning to saturate above the level of species. This is consistent with a recent extensive faecal microbiome survey of goats and sheep that found only 3.7% novelty at the genus level for their MAG dataset, but 84.5% novelty at the species level ^47^.

Read mapping analysis against the 3,868 marsupial MAG dataset and 85,205 GTDB 09-RS220 species representatives (*see Methods*) ^46^ revealed that a substantial amount of microbial diversity was not captured in the MAG or reference genome sequences for most marsupial hosts with a median read mapping percentage of 40% (ranging from 5 to 70%; **Fig. S4**). This is comparable to a previous animal faecal microbiome study in which a large proportion of metagenomic reads did not map to reference genomes (21%) ^48^ suggesting that total diversity in the gut is still far from saturation. Consistent with this inference, a marker gene analysis of the marsupial scat reads using SingleM indicates substantially more prokaryotic microbial diversity remains to be described including 2 novel phyla, 27 novel classes, 59 novel orders and 132 novel families (**Table S19**). Differences in read recruitment observed between marsupial hosts partly reflect prior metagenomic studies, with the koala and wombat ^9,10,28^ having amongst the highest levels of read recruitment (**Fig. S4**).

### Broad marsupial host range species

Only 10.8% of marsupial gut prokaryotic species represented by genome sequences were found in two or more marsupial families (**Table S20**). The most ubiquitous species in our study, found in 35 individual scat samples across all six marsupial families was *Escherichia coli*. This well-studied microbial species is a facultative anaerobe and common in the lower intestine of mammals, although it typically constitutes only a small percentage of the total prokaryotic community. *E. coli* is also found widely in the environment ^49^ suggesting that it likely moves between marsupial hosts, which is supported by its presence in cohabiting individuals at Port Douglas Wildlife Habitat and low incidence at other animal sanctuaries (**Table S20**). Other broad range species include *Phascolarctobacterium faecium* and *Akkermansia muciniphila* detected in five of the six marsupial families investigated (**Table S20**). *P. faecium* was originally isolated from koala faeces ^50^, but has since been found in a wide range of animal hosts including human ^51^, rat ^52^ and cow ^53^. It uses succinate as a primary substrate, likely to be provided by *Bacteroides thetaiotaomicron* ^54^ or other *Bacteroides* species that are also commonly found in marsupials (**Table S20**). *A. muciniphila* is a specialist mucin degrader common to many animal species ^55^ and is likely to play the same role in marsupials.

### Marsupial host specific taxa

The majority of microbial species with genomic representation were lineage specific (89.2%; **Table S20**) consistent with separation of host marsupial families by their taxonomic profiles (**Fig. 2A & B**). *Cryptobacteroides* species are conspicuous constituents of marsupial grazers particularly in kangaroos, which harbour 48 novel species (**Table S20**) including a radiation of 37 species indicating a macropod-specific expansion of this genus (**Fig. 3 & 4A**). *Cryptobacteroides* are recognised for their importance in the rumen where they are inferred to degrade lignocellulose and starch via polysaccharide utilisation loci (PUL) ^56^ and may play a similar role in kangaroos. Relative to other marsupial *Bacteroidota,* the *Cryptobacteroides* species are enriched in starch-, xylan-, β-glucan- and β-mannan-degrading enzymes, indicating their likely importance in grass digestion in kangaroos and wombats (**Table S21**).

**Figure 3.**
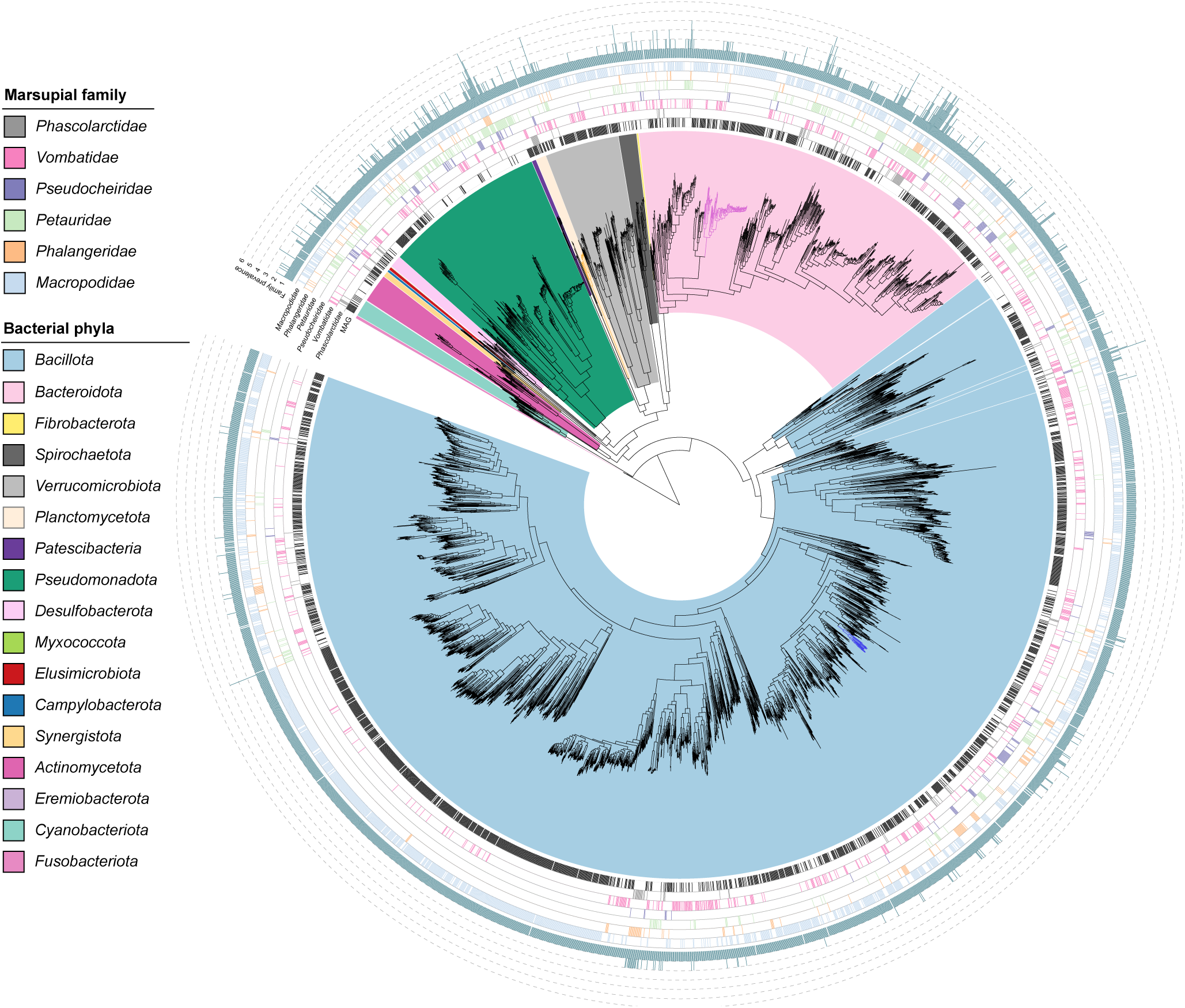
Maximum-likelihood phylogenetic tree of dereplicated bacterial MAGs supplemented with public genomes identified as present in marsupials based on read mapping to the GTDB or profiling with SingleM. Rings indicate (from inside): MAGs from the current study, presence in marsupial families based on read mapping and count of marsupial families bacterial species is present in. Dark orange branches indicate the class *SZUA-567*, dark pink the genus *Cryptobacteroides* and dark blue the genus *Eubacterium_I* (see Fig. 4 for more detail).

Consistent with a previous report ^9^, an as-yet-uncultured basal *Planctomycetota* lineage, *SZUA-567*, was identified in koala scat. This family-level lineage is represented by two genera and nine species in our survey and was found solely in koalas (**Table S20**), suggesting that it was introduced into the koala microbiome after divergence from wombats approximately 35-40 Mya ^57^ (**Fig. 1**). Currently, the closest relatives of koala *SZUA-567* MAGs have been identified in termite gut microbiomes ^58^. Koalas are known to feed on termite nests ^59^ and *SZUA-567* species from koalas and insects are interleaved (**Fig. 4B**) leading us to hypothesise that ingested termites are the source of this lineage in the koala. The specific role of these bacteria and how they managed to colonise the koala gut remain to be determined.

**Figure 4.**
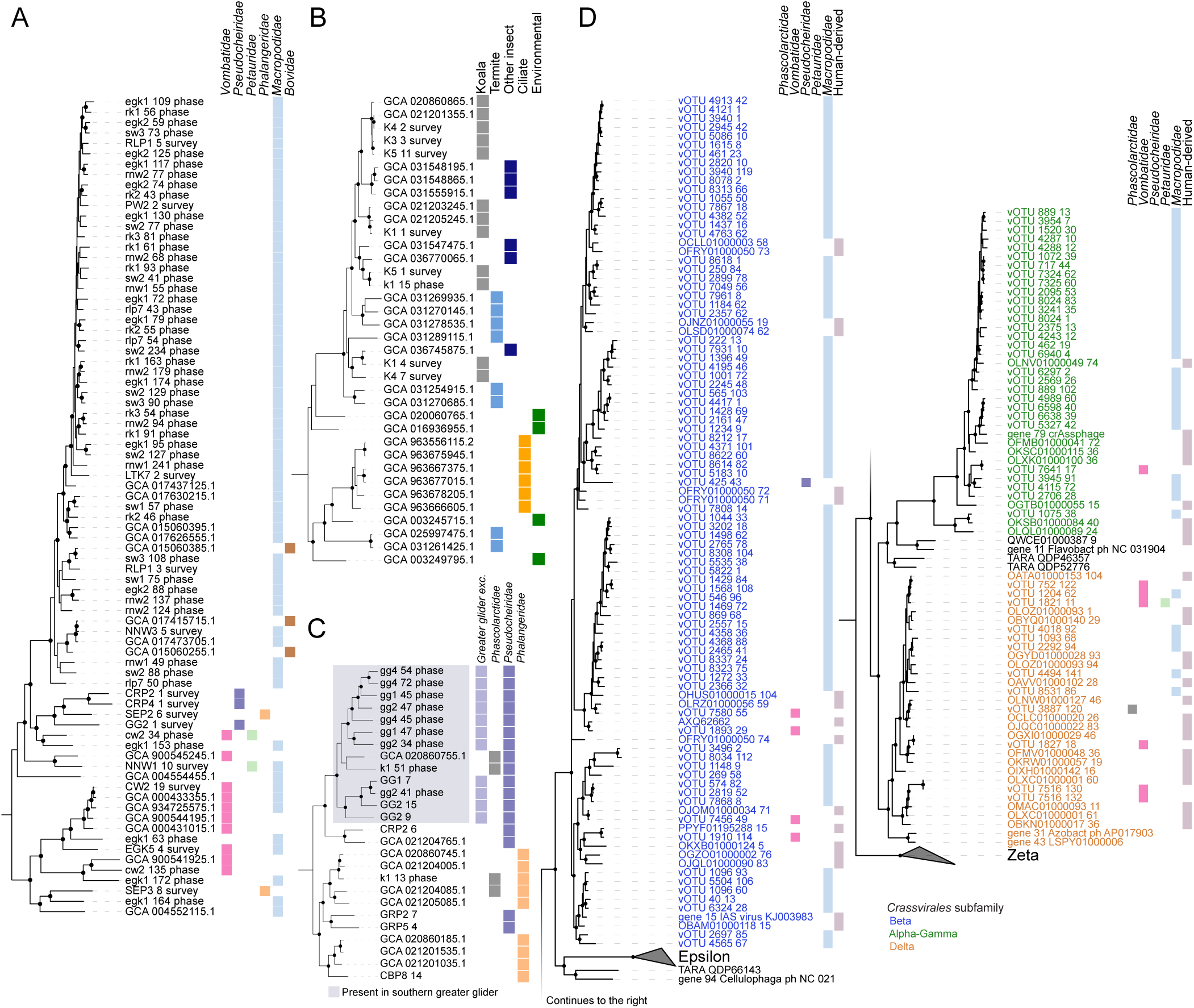
Maximum-likelihood phylogenetic trees of **a**, *Cryptobacteroides* **b**, family *SZUA-567* **c**, *Eubacterium_I* and **d**, *Crassvirales*. Trees in panels **a-c** incorporate dereplicated bacterial MAGs and public genomes and are inferred from an alignment of GTDB marker proteins ^46^. Panel **d** tree incorporates vOTU representatives and public *Crassvirales* genomes used in ^81^ and is based on the alignment of TerL proteins. Boxes to the right of each tree indicate presence of each species in marsupial families based on read mapping or origin of public genomes.

An expansion of the *Lachnospiraceae* genus *Eubacterium_I* was identified in southern greater gliders with 13 species identified, of which 11 were found exclusively in this host (**Fig. 4C**) and the other two also found in koala scat (**Table S20)** ^28^. The only characterised representative of this genus, *E. ramulus* isolated from human faeces, is recognised for its ability to degrade flavonoids ^60,61^. Since flavonoids are prominent phenolic metabolites in eucalyptus leaves including catechins, isorhamnetin, phloretin, kaempferol and luteolin ^62,63^, we hypothesize that the role of *Eubacterium_I* in the greater glider and koala is to metabolise these compounds. This is supported by enrichment of flavonoid modifying enzymes in marsupial *Eubacterium_I* species relative to other marsupial *Lachnospiraceae*, including flavone/flavonol reductase (*flr*, which initiates flavone and flavonol catabolism in the gut ^64^, flavanone/flavanol-cleaving reductase (*fcr*) and chalcone isomerase (*chi*; **Table S22**).

### DNA viruses of the marsupial gut

To assess marsupial gut viromes, we focused our analysis on the subset of 23 host samples sequenced using proximity ligation due to greater sequencing depth (**Table S3**) and the ability to connect non-integrated viruses to their hosts. A total of 12,142 DNA viral genomes were obtained ranging in size from 18 to 231 kbp (average 48.6 kbp) (**Table S23**). These represent 8,928 viral species (vOTUs) according to a recommended ANI-based definition ^65^ (**Table S23**). Most of these species were represented by a single genome (82%), however 21 vOTUs comprised 10 or more genomes each (**Table S23**). The great majority (99%) of the vOTUs had no close (species-level) matches in public databases (**Table S24**). However, at a higher taxonomic level, 98% of the marsupial viral species belong to the class *Caudoviricetes* (**Table S23**) consistent with DNA viruses found in many other animal faecal microbiomes ^66–68^ and morphological identification of a great variety of *Caudoviricetes* (*Myoviridae*, *Siphoviridae* and *Podoviridae*) in the gastrointestinal tracts of kangaroos ^69^.

### Prokaryote and marsupial host range

Almost a third of the marsupial vOTUs (2,713) could be linked to one or more prokaryotic host species either via proximity ligation (as evidenced by interlinkage reads) and/or by host genome sequences flanking the viral genome (**Table S25**). Most of these vOTUs (2,405; 89%) were associated with a single host species (**Table S25**) consistent with the typically narrow host range of cultured phages ^70^. Further, culture-independent evidence from human faecal samples using single cell viral tagging found that 98% of the phages targeted only one host species ^71^. Together these findings suggest that phages are not significant vectors of gene exchange between different bacterial species in the marsupial gut microbiome and animal gut more generally. Of the 308 vOTUs with links to >1 prokaryotic host species, 42% (129) were genus specific, 31% (96) family specific, 16% (51) order specific, 8% (25) class specific, 2% (5) phylum specific and 1% (2) domain specific (**Table S25**). The most promiscuous viral species, vOTU_2a, was linked to nine host species belonging to five genera of the family *CAG-272* in the order *Oscillospirales* (**Table S25**). Another viral species, vOTU_14, appears to be capable of targeting monoderms (a member of the spore-forming *Bacillales* genus *Pradoshia*) and diderms (*Prevotella sp017410365*), a rarely observed trait amongst phages ^72^. The carriage of multiple genes encoding phage tail lysozyme domains in vOTU_14 genomes may assist this virus in breaching different cell envelope types (**Table S26)** ^73^.

The marsupial host range of faecal viral species reflected in large part prokaryotic populations in that most were restricted to single host lineages; 59% (5,282) to a single marsupial host species, 67% (6,001) to a single genus and 96% (8,584) to a single family (**Table S27**). Indeed, a substantial portion of vOTUs (3,692, 41.4%) were identified in a single faecal sample, suggesting that many are specific to individual animals, consistent with the observed individuality of human gut viral communities ^74,75^ (**Table S27**). Some of these localised viral populations were linked to bacterial species found in multiple marsupial families (e.g. vOTU_1974 to *Bacteroides xylanisolvens* and vOTU_7758 to *Parabacteroides distasonis*; **Tables S20 & S25**) suggesting that they have not been transferred between marsupial species via their prokaryotic hosts. This is similar to an observation in human faeces where the same bacterial species was targeted by different viruses in different faecal samples ^71^. The converse was also found, however, where the broad marsupial host range of a bacterial host was mirrored by their viruses. For example, the only viral species identified in representatives of five marsupial families (vOTU_1924; **Table S27**) was linked to *Phascolarctobacterium faecium* detected in the same marsupial families (**Table S20**), suggesting that this virus has been successfully dispersed between marsupial species with its host.

### Crassvirales are prevalent in macropods

*Crassvirales* are a recently discovered lineage of dsDNA viruses that have gained attention due to their prevalence in the faecal microbiomes of humans and other primates ^76–78^. However, their presence in other animal hosts appears to be sporadic, with the exception of chickens ^79^. Notably, 151 (1.7%) of the marsupial vOTUs were classified as *Crassvirales*, most of which (91%, 137 vOTUs) belong to macropod clades (**Fig. 4D**; **Table S27**). Consistent with previous reports of a strictly lytic lifestyle, marsupial *Crassvirales* were not found to be integrated into their prokaryotic hosts (with one potential exception; rk1_vMAG_90 was predicted to be a provirus, however, no host MAG was identified), and lack identifiable integrases and recombinases necessary for lysogenic infection (**Table S26**) ^80^. Only five *Crassvirales* genomes were linked to identifiable host MAGs, three members of the *Bacteroidota* (in the genera *Alistipes*, *Barnesiella* and *Prevotella*), and two *Bacillota* (both members of the genus *Candidatus* Coprovivens). This appears to extend the known host range of these viruses, which are thought to exclusively target members of the phylum *Bacteroidota* ^78,79^, although CRISPR spacers suggest broader associations, including with members of the *Bacillota* ^81^. *Ca.* Coprovivens belongs to the as-yet-uncultured *Bacilli* order *RF39*, all members of which are predicted to be host-associated ^82^. Therefore, it is possible that *Bacteroidota* cells simply co-host *Ca*. Coprovivens and *Crassvirales*.

### Antimicrobial resistance potential in marsupials

As captive animals comprise a large portion of marsupials in this study, we assessed the antimicrobial resistance (AMR) potential of their scat microbiomes across the dataset to understand whether this was influenced by captivity as is seen in other species species ^83,84^. We identified 188 types of recognised AMR genes across the 117 marsupial scat samples (3,907 total genes), representing 24 classes of antimicrobial resistance genes (**Table S28**). Approximately half (47%) of these gene types were identified on potentially mobilisable elements (plasmid or viral contigs), with plasmids carrying ∼20 to 40-fold more AMR genes (1 in ∼1,500 genes) than viral (1 in ∼60,000 genes) or chromosomal sequences (1 in ∼33,000 genes; **Fig. S5**), consistent with the primary role of plasmids in AMR transfer ^85^. Plasmid-encoded AMR genes were also more diverse than viral-encoded genes, comprising 17 AMR classes *vs* 10 for viral sequences (**Table S28**).

The most prevalent classes of resistance genes found in marsupial faeces target glycopeptides (93% of captive animals, 86% wild) and ꞵ-lactams (87% captive, 45% wild) (**Table S29**), broadly reflective of the widespread distribution of these AMR classes ^86^. Both of these classes were significantly enriched in captive *vs* wild animals, as were lower prevalence classes targeting macrolides (80% vs 5%), lincosamides (75% vs 5%), sulfonamides (28% vs 0%) and trimethoprims (20% vs 0%) (**Table S29**). While captive animals harboured a significantly higher abundance of many AMR classes overall, this effect was host-dependent (**Fig. S6**, **Table S29**). Captive possums, southern greater gliders and koalas (families *Phalangeridae*, *Pseudocheiridae* and *Phascolarctidae*) typically displayed lower AMR gene abundance than macropods, wombats and small gliding species (families *Macropodidae*, *Vombatidae* and *Petauridae*), with the exception of one hospitalised common brushtail possum (CBP3) (**Table S30**). Human intervention via antibiotic treatment is therefore likely driving differences in prevalence of AMR gene classes between captive and wild animals, and between different marsupial families. For example, in macropods, clindamycin (a lincosamide) is used to treat necrobacillosis and co-trimoxazole (a combination of sulfonamide and trimethoprim) to treat wounds and urinary tract infections ^87,88^. However, other antibiotics such as macrolides are not commonly used to treat marsupials ^89^ and increased macrolide resistance in captive animals may be due to other factors, such as transfer of antibiotic resistant bacteria from humans and other animals or environmental sources ^90,91^.

### Concluding remarks

In this study, we present a metagenomic (including proximity ligation data) and metabolomic molecular inventory of the faecal microbiomes of 23 marsupial species, the first such data for 13 of these species, and the first metabolomic and proximity ligation data for any marsupial. The 3,868 assembled medium-to-high quality MAGs include representative genomes for 1,951 prokaryotic species (80%) and 58 genera (9%) new to GTDB 09-RS220. In addition, we present a marsupial viral genome catalogue of 12,142 genomes (8,928 vOTUs), with prokaryote host links for ∼30% of viral species. Combined, these data confirm host family as the primary driver of marsupial faecal taxonomic, functional and metabolomic profiles. Host specificity extends to species with highly specialised, eucalyptus-dominant diets (koalas and southern greater gliders), where their distinct taxonomic and functional profiles support alternative solutions to eucalyptus digestion. We observe host lineage-specific bacterial expansions including *Cryptobacteroides* species in kangaroos, the *Planctomycetota* family *SZUA-567* in koalas and *Eubacterium_I* in greater gliders and expanded repertoire of *Crassvirales* phage in macropod hosts. We expect these data will provide a valuable reference for comparative analyses of other animal faecal and gut microbiome datasets. While this resource expands our knowledge of the marsupial microbiome, many species remain entirely unexplored (122/151 *Diprotodontia* species), indicating the need for ongoing work in this area.

## Methods

### Sample collection

Ninety-five faecal samples from 82 captive marsupials were collected from wildlife sanctuaries in South-East Queensland (Lone Pine Koala Sanctuary, Brisbane; David Fleay Wildlife Park, Burleigh Heads; and Currumbin Wildlife Sanctuary, Currumbin) and North Queensland (Wildlife Habitat, Port Douglas and Cairns Tropical Zoo) (**Table S3**). Twenty-two samples from 16 wild marsupials were collected from areas surrounding Brisbane and Wongabel State Forest, Queensland. Samples were stored in Eppendorf tubes at −80°C prior to downstream processing.

### DNA extraction and sequencing of Illumina-only samples

For the 94 faecal samples sequenced using only Illumina sequencing, DNA extraction and sequencing methods are described in ^28^. Briefly, ∼50 mg of faecal sample was added to tubes with 0.7 mm garnet beads (MO BIO, Carlsbad, CA, USA) and suspended in 750 µL Tissue Lysis Buffer (Promega, Madison, WI, USA). Following 10 min of bead-beating at maximum speed, samples were pelleted by centrifugation. Finally, 200 µL of supernatant was used as the input for DNA extraction on a Maxwell 16 Research Instrument with the Maxwell 16 Tissue DNA Purification Kit (Promega, Madison, WI, USA) according to the manufacturer’s instructions. DNA was sequenced with the Nextera XT DNA Sample Prep kit (Illumina, San Diego, CA, USA). Samples were sequenced on the Illumina NextSeq 2000 platform, producing 150 bp paired-end reads at the Australian Centre for Ecogenomics.

### Proximity ligation sample preparation

For the 23 samples sequenced by Phase Genomics using proximity ligation sequencing, faecal pellets were cut in half with a sterile scalpel blade in a laminar flow hood and the centre of the pellet was added to a sterile tube with PG Shield (Phase Genomics, Seattle, WA, USA). For southern greater glider samples, due to the limited amount of available sample, ∼0.5 g of faeces was added to 5 mL of PG Shield. For the remainder of the samples, ∼1 g of faeces was added to 10 mL of PG Shield. Tubes were shaken and sent to Phase Genomics for proximity ligation sequencing using the ProxiMeta Hi-C v4.0 Kit which sequentially cross-links cells, mechanically lyses them, isolates and fragments chromatin, and ligates spatially adjacent DNA ends. After reversing cross-links, the biotin-tagged Hi-C junctions were purified on streptavidin beads to yield DNA ready for paired-end sequencing ^92^. Indexed Hi-C libraries were pooled and loaded onto an Illumina NovaSeq X with 2 × 150 bp chemistry.

### Metagenome-assembled genome recovery

For the Illumina-only samples, sequencing adaptors were removed with SeqPurge (v2018_11) ^93^ and the trimmed reads were assembled into contigs using MetaSpades (v3.15.3) using default settings ^94^. The contigs were binned using Aviary (v.0.4.3) with aviary recover in default mode ^95^ from coverage profiles generated across all samples within each main animal group (kangaroo/wallaby, possum, koala, wombat).

For the proximity ligation samples, sequencing reads were processed as described ^37^, with assembly performed using MEGAHIT (v1.2.9) ^96^ and binning performed using the ProxiMeta platform ^97,98^ incorporating both standard shotgun and Hi-C data.

### Marsupial MAG database

MAG completeness and contamination was assessed using CheckM2 (v.1.0.1) ^99^. MAGs with a completeness of ≥70%, contamination of ≤5% and an N50 of ≥10 kb were chosen for downstream analysis supplemented with publicly available genomes. Public genomes for inclusion in the genome database were selected from species representative genomes in GTDB 08-RS214 ^46,100^. Selection was based on read mapping values generated using CoverM v0.6.1 ^101^, filtering for alignments exceeding 95% identity across 90% of the read length. Public genomes achieving ≥0.05% relative abundance and recruiting reads to ≥10% of the genome length were included in the final database (n=1,201). The combined genome set (3,868 MAGs + 1,201 public genomes) was dereplicated using CoverM v0.7.0 ^101^ with the ‘cluster’ command at 95% identity with an alignment fraction of 60%. The ‘Parks2020_reduced’ quality formula was used with genome statistics generated by CheckM2 v1.0.1 ^99^. MAGs were classified with GTDB-Tk using GTDB release 09-RS220 ^102^. A maximum likelihood tree was inferred using IQ-TREE v2.2.2.3 ^103^ based on the alignment generated by GTDB-Tk ‘de_novo_wf’, using ModelFinder for model selection within the LG model set. Ultrafast bootstrap approximation was generated from 10,000 replicates.

### Marsupial host DNA contamination

As there were no genomes available for the majority of species in the dataset (genomes currently available for koala, eastern-grey kangaroo, yellow-bellied glider, common ringtail possum, green ringtail possum, common brushtail possum & common wombat), an estimate of host DNA contamination was established via read mapping to a marsupial genome database comprising available genomes and exome sequencing data from all genera in the dataset (**Fig. S7** ^104–106^, **Table S31**). Read mapping was performed using CoverM v0.7.0 ^101^ filtering for alignments exceeding 95% identity across 90% of the read length.

### Marsupial host tree

Nine genes (*apoB*, *brca1*, *irbp*, *vwf*, *cytb*, *coi*, *nadh2*, 12S rRNA, 16S rRNA) from the 23 marsupial species used in this study were searched for in the NCBI database (**Table S32**) ^107^. For the three marsupial species that were missing from the NCBI database, the missing genes were identified from the sample metagenome assemblies using BLASTx (v.2.9.0) ^108^. The BLAST hit with the highest e-value was extracted from the assemblies using BEDTools (v2.27.1) ^109^ and the correct open reading frame was identified by translating the nucleotides to amino acids using https://web.expasy.org/translate/. An NCBI BLASTp search was conducted to determine if the correct gene had been identified. MAFFT (v7.505) ^110^ in auto mode was used to align the genes. The alignment was trimmed with trimAl (v1.4.1) ^111^ using the gappyout mode. A phylogenetic tree of the multiple sequence alignment was created using IQ-TREE (v1.6.12) ^103^ using ModelFinder automatic model selection to identify the best-fit model and 1,000 bootstrap replicates. The concatenated gene tree was visualised in ARB (v6.0.6) ^112^ and rooted on the outgroup (*Dromiciops gliroides*) before being exported into Adobe Illustrator 2023.

### Prokaryotic community composition

Reads were deduplicated using BBMap v39.01 (sourceforge.net/projects/bbmap/) (dedup.sh; absorbrc=f absorbmatch=t absorbcontainment=f) and quality trimmed using Trimmomatic v0.39 ^113^ (SLIDINGWINDOW:4:15 LEADING:3 TRAILING:3 MINLEN:50) prior to mapping to the compiled genome database (described above). Deduplicated, untrimmed reads were used for marker gene-based community analysis. Illumina reads generated by Phase Genomics were subsampled to 10% of their full depth using SeqKit v2.4.0 ^114^ to align with the read depth of other samples.

Genome-based community analysis was performed using relative abundance values generated from read mapping using CoverM v0.7.0 ^101^. Read alignments were filtered to retain the subset that had ≥95% identity across ≥90% of the read length, excluding alignments to genomes with a minimum covered fraction of ≤10%. Marker gene-based community analysis was based on read classification using SingleM v0.16.0 ^115^. Per-marker counts were combined using the ‘condense’ function. The bacterial and archaeal community fraction was estimated using SingleM Microbial Fraction ^116^. Shannon diversity was calculated from SingleM coverage values using phyloseq v1.48.0 ^117^.

### Functional profiling

Proteins within assembled contigs were identified using Prodigal v2.6.3 ^118^ and clustered at 90% identity across 80% of protein length using MMseqs2 v14-7e284 ^119^. Representative protein sequences were functionally annotated using DRAM v1.5.0 ^120^ using the Kofam ^121^, Pfam ^122^ and dbCAN ^123^ databases accessed December 2022.

Sample reads (forward only) were aligned to the protein database using DIAMOND v2.1.0 ^124^ (settings evalue 0.00001, min-score 40, query-cover 80, id 70, max-hsps 1 and max-target-seqs 1), filtering for reads ≥ 140 bp with Seqkit v2.4.0 ^114^. Reads were also aligned to the single copy marker genes included within the SingleM GTDB release 08-RS214 metapackage ^115^ using the same settings. RPKM (reads per kilobase per million mapped read) values were calculated for both the protein database and SingleM marker genes, with the mean RPKM value across all SingleM markers used to normalise protein database values on a per sample basis. Normalised RPKM values per protein were summed across equivalent functional annotations and log_2_ transformed for PCA and differential abundance analysis.

### Virus identification, filtering and taxonomic assignment

Viral contigs in the proximity ligation assemblies were identified using VIBRANT (v1.2.1) ^125^, with contigs carrying both bacterial and viral sequence annotated as prophage where viral sequence comprised ≥50% of the contig length. Viral binning was performed using the ProxiPhage algorithm as described ^37^. Briefly viral contigs are clustered into viral MAGs (vMAGs) by merging proximity-ligation signal–based clusters and conventional binning results via a greedy network-collapse algorithm, yielding non-redundant vMAGs that meet minimum Hi-C linkage thresholds. Putative viral hosts were identified based on Hi-C linkage data ^37^. To limit false positive interactions, only virus-host linkages supported by ≥2 Hi-C read links, a connectivity ratio of ≥0.1 and copy number meeting a threshold determined by a ROC curve were used ^37^. Connections with an average copy count of less than 80% of the highest copy count were also removed.

To minimise analysis of short viral fragments, we set a viral genome length threshold of ≥18 kb, which is the recommended minimum length of a member of the *Caudoviricetes* ^126^, the dominant viral class found in gut microbiomes ^66–68^. GeNomad v.1.8.1 ^127^ was used to assign taxonomy to the viruses using the end-to-end command in default mode. For the vMAGs (viruses with multiple contigs), the contigs were concatenated before running GeNomad. Viral novelty was assessed via BLAST alignment to the NCBI viral catalogue (downloaded August 2024) and the Unified Human Gut Virome Catalog (UHGV, votus_full dataset, accessed February 2025, https://github.com/snayfach/UHGV), retaining hits with ≥95% identity and ≥80% alignment fraction (of query sequence).

### Crassvirales validation

Candidate terminase (TerL), major capsid protein (MCP) and portal proteins were identified in *Crassvirales* genomes using HMM profiles generated from published multi-sequence alignments ^128^ using hmmer v3.3.2 ^129^. Marsupial proteins of length ≥500 with HMM alignment E-value ≤1e-10 (TerL, portal) or 1e-06 (MCP) were combined with known *Crassvirales* and outgroup proteins of each type ^81^. Sequences were aligned using Mafft v7.490 ^110^ and alignments trimmed using trimAl v1.5.0 ^111^ with the ‘gappyout’ option. Maximum-likelihood trees were inferred using IQ-TREE v2.4.0 ^103^ with the ModelFinder option and bootstrap support approximated from 10,000 ultrafast bootstrap replicates.

### Viral clustering and community profiling

An initial all-vs-all comparison of viral genomes was undertaken using skani (v0.2.2) ^130^. The Marker k-mer compression factor parameter (-m) was set to 200 (recommended for viruses), and the --slow preset was applied to improve average nucleotide identity (ANI) accuracy, particularly for highly fragmented assemblies. Genome pairs with ANI ≥ 95% and alignment fraction (AF) ≥ 85% were selected, following the thresholds used in ^131^. The selected genome pairs were combined into clusters of viruses with similar ANI. Each cluster was manually inspected to identify and resolve non-robust connections. For each cluster, a representative genome was selected based on its connectivity within the cluster.

The genome with the highest number of ANI connections was chosen as the representative. If multiple genomes had the same maximum number of connections, the genome with the highest average ANI across all its connections was selected as the representative.

The viral community across all samples was assessed based on a viral database containing all cluster representatives and singleton genomes. Read mapping of all samples to the compiled database was performed using CoverM v0.7.0 ^101^, filtering alignments ≥95% identity across ≥90% of the read length, with a minimum covered fraction of 30%.

### Antimicrobial resistance

Potential antimicrobial resistance genes were identified across all assembled contigs with AMRFinderPlus (v3.12.8) ^132^ on proteins identified using Prodigal (v2.6.3) ^118^. Hits were filtered for those ≥50% identity across ≥80% protein length. Sequence source of each AMR gene (viral, plasmid or chromosomal) was based on prediction of plasmid/viral sequences for each contig using GeNomad v1.8.1 ^127^. AMR class abundance was calculated as the sum of RPKM values protein within that class, generated as described above. Differential abundance of AMR classes between wild and captive animals was performed using MaAsLin2 ^133^ with marsupial species as a random effect. Marsupial family comparisons (captive animals only) included location as a random effect. Data were log_2_ transformed prior to analysis.

### Metabolomic profiling

Metabolites were extracted from faecal and plant samples using hexane according to ^134^. In brief, ∼100 mg of frozen homogenized material was added to 1 mL of hexane, followed by incubation (1 h, 40°C, gentle agitation). Thereafter debris was precipitated by centrifugation (5 min, 2000 x g), and dried overnight (65°C) to obtain dry mass for normalization. The resultant supernatant (1 μL) was analyzed by GC–MS (GCMS-QP 2010 Plus, Shimadzu) using a stationary phase column (HP-5 ms Ultra Inert GC Column, 20 m x 0.18 mm, 0.18 μm, 7 inch cage, Agilent Technologies), with a purge flow of 4 mL min−1 and hydrogen as carrier gas. Injection temperature was 250 °C in splitless mode with a temperature gradient of: 40 °C, 4 min hold; 2 °C min−1 to 120 °C; 30 °C min−1 to 280 °C; 3 min hold at 280 °C. Obtained chromatograms were analyzed using PARADISe ^135^.

Chromatograms were processed using PARAFAC2-based PARADISe dashboard ^135^ for simultaneous deconvolution and integration. The tentative identity of individual unknowns was accepted by comparison of their mass-spectral data with those of the NIST11 mass spectral library (NIST) data after comprehensive peak alignment, manual interval selections, and feature detections. Further processing of peak areas was performed using the web portal of MetaboAnalyst (v5.0) ^136^. Missing values were replaced with 20% of the minimum value for that compound and compounds with ≥50% missing values were removed. Compounds with low repeatability (relative standard deviation ≥30%) and low variance (10% of total compounds) were also removed. Remaining data were median normalized, log-transformed and auto-scaled prior to analysis in R.

### Statistical methods and data visualisation

SingleM coverage and read mapping-based data (genome-based prokaryotic and viral profiles) in relative abundance format were centred log-ratio transformed prior to PCA and differential abundance analysis. Prokaryotic genome-based profiles were filtered for taxa present in ≥2 samples. Squared Shannon diversity values were compared using lme4 v1.1-35.4 ^137^ and lmerTest 3.1-3 ^138^ with marsupial species as a random effect. RPKM data (KEGG, CAZy, AMR profiles) were log_2_ transformed prior to analysis. Metabolomic data preparation is described above. Where multiple samples were obtained from the same animal, only a single sample was retained for statistical analyses.

PCA and redundancy analyses were conducted using vegan v2.6-4 (R v4.3.2) ^139^. To identify the most influential features in PCA, species scores were extracted and the Euclidean magnitude of each feature across PC1 and PC2 was calculated, and the top 20 features by magnitude were selected for display. Redundancy analysis included marsupial family, diet (grazer, browser, omnivore, fungivore), gut type (hindgut, foregut), location and captivity status as explanatory variables and significance was determined using the ‘anova.cca’ function. Enrichment of functions between groups was assessed using EnrichM v0.6.1 (https://github.com/geronimp/enrichM). Plots were created using ggplot2 v3.5.1 ^140^. Bacterial and viral phylogenetic tree figures were created using iTOL ^141^.

## Author contributions

RMS, MM, BLM, EHJN and PH designed the project. RMS, MS, DG, MM, MI, IL, BA and EHJN performed experimental work. RMS, KLB, PAC, DLAW, BA, LK and EHJN performed data analysis. JZ provided data analysis scripts. All authors contributed to data interpretation. RMS, KLB and PH wrote the manuscript and all authors edited and approved the submitted version. RMS and KLB contributed equally to this work.

## Conflicts of interest

IL and BA are employees of Phase Genomics, Inc.

## Funding information

This work was supported by Australian Research Council Discovery Projects DP150104202, awarded to P.H. and M.M, and DP250103673, awarded to P.H. and K.L.B, and partially supported by grant INV-044643 from the Gates Foundation to Phase Genomics.

## Ethical approval

Ethical permission for the collection of all samples was granted by the Animal Ethics Unit, the University of Queensland, Brisbane, Australia, under ANRFA/SCMB/099/14 and 2023/AE000378.

## Acknowledgements

We thank Lone Pine Koala Sanctuary, David Fleay Wildlife Park, Wildlife Habitat Port Douglas, and Cairns Tropical Zoo for providing samples from captive animals and D.S. Teakle and A. Shima for the collection of samples from wild animals. We thank Miriam Shiffman and Emily Hoedt for sample collection and faecal DNA extractions, Alex Hasson for preliminary statistical analysis of the data and Thomas Sicheritz-Pontén for discussions.

---

**Fig. S1.**
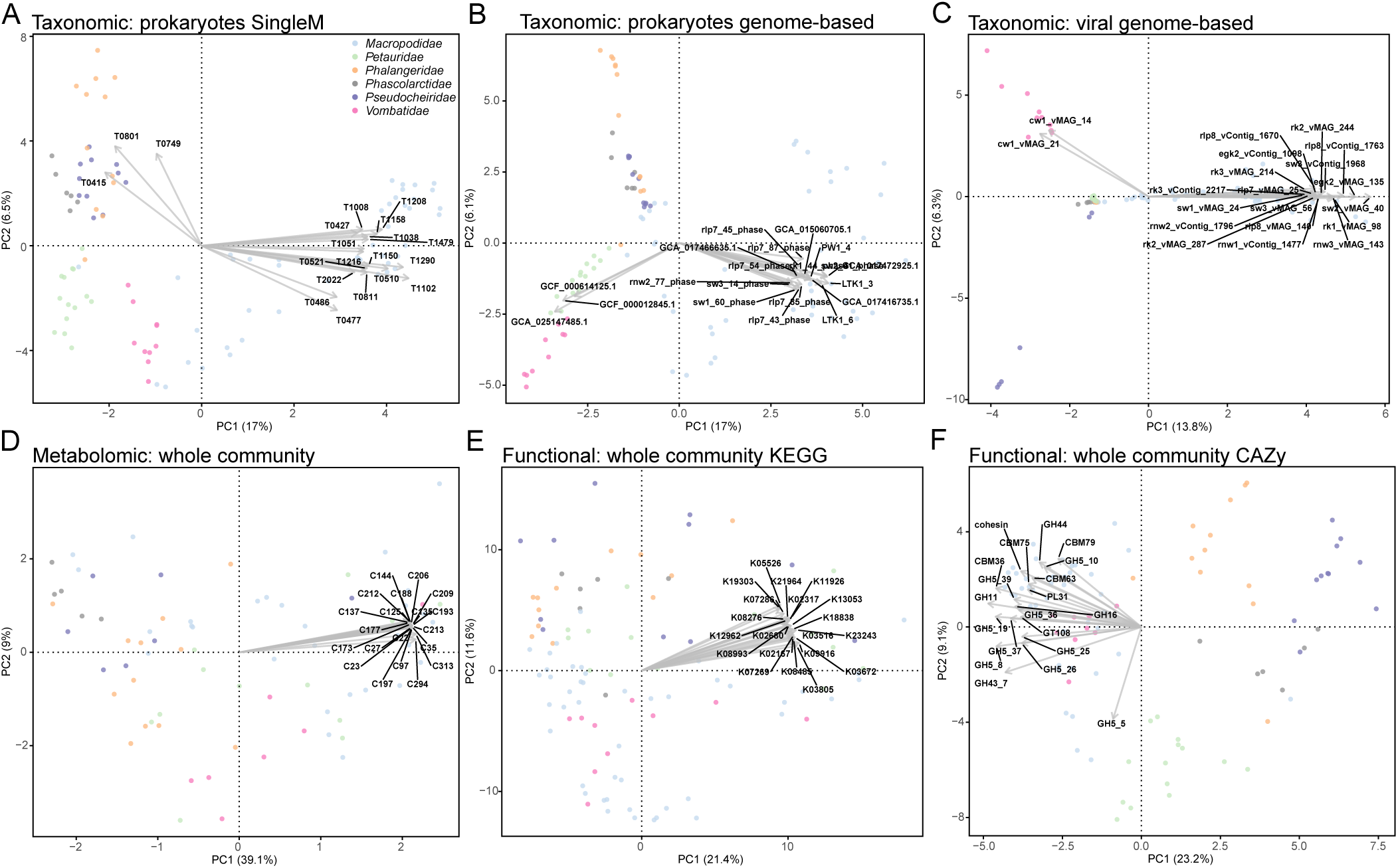
Principal component analysis based on **a**, marker gene-based prokaryotic community **b**, genome-based prokaryotic community **c**, genome-based viral community **d**, metabolomic **e**, KEGG-based functional and **f**, CAZy-based functional profiles. Top 20 features based on Euclidean magnitude across PC1 and PC2 are displayed. Full lists are contained in Tables S4-S9.

**Fig. S2.**
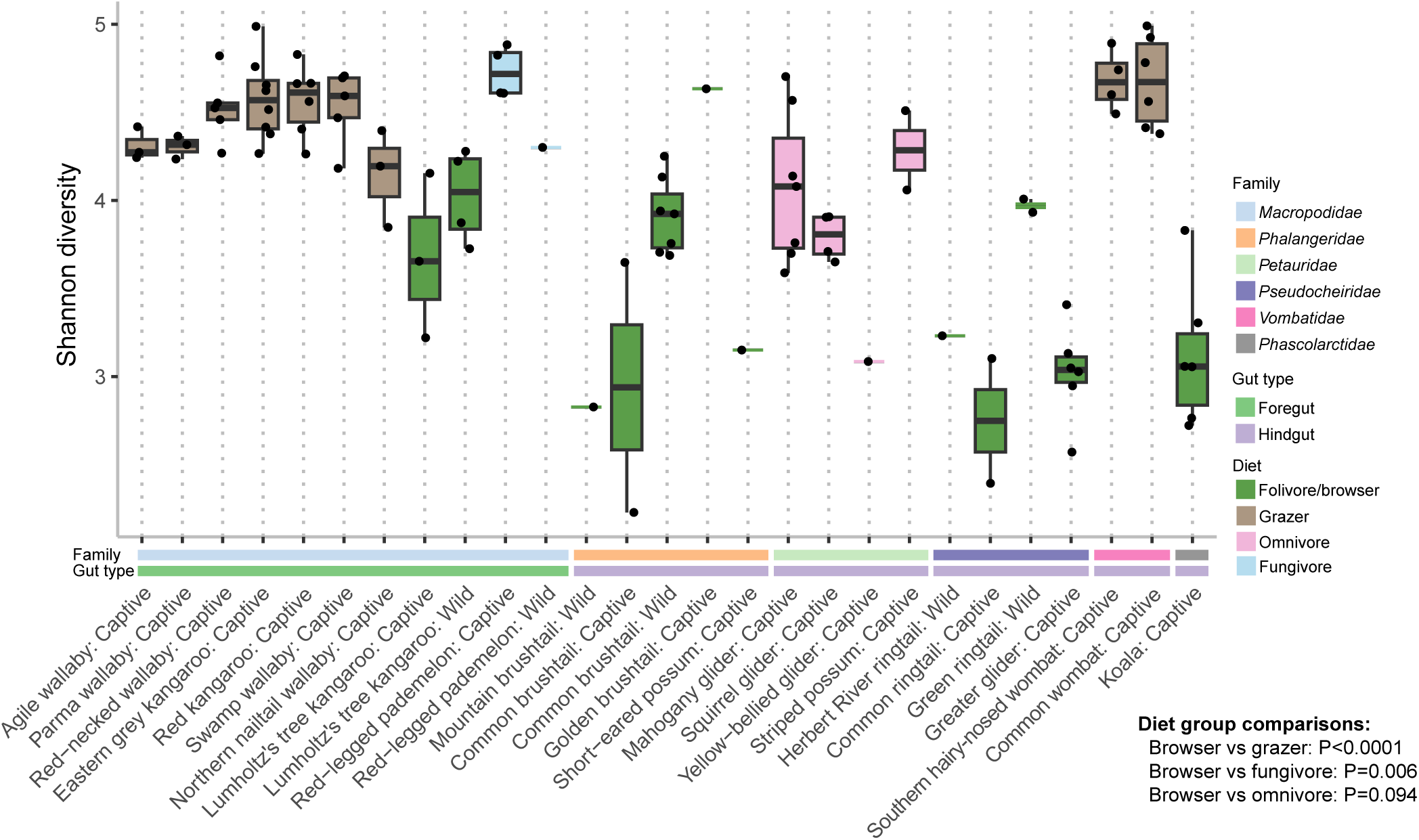
Shannon diversity based on SingleM prokaryotic community profiles.

**Fig. S3.**
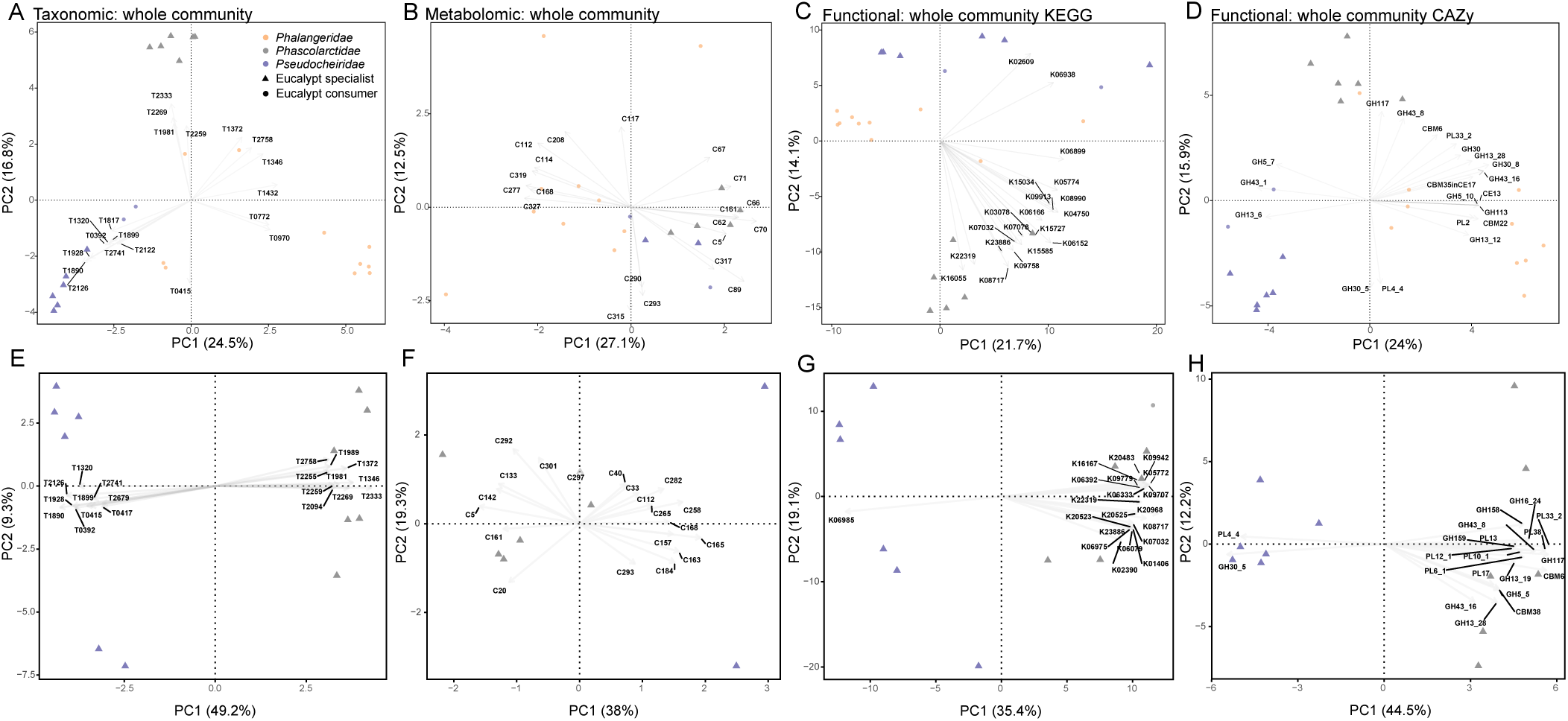
Principal component analysis of eucalypt consuming marsupials (**a-d**) and eucalypt specialist marsupials (**e-h**) based on a & e, marker gene-based prokaryotic community, **b & f**, metabolomic, **c & e**, KEGG-based functional, **d & h**, CAZy-based functional profiles. Eucalypt consumers includes koalas, greater gliders, common and golden brushtails and common ringtails. Eucalypt specialists include koalas and greater gliders. Top 20 features based on Euclidean magnitude across PC1 and PC2 are displayed. Full lists are contained in Tables S10-S17.

**Fig. S4.**
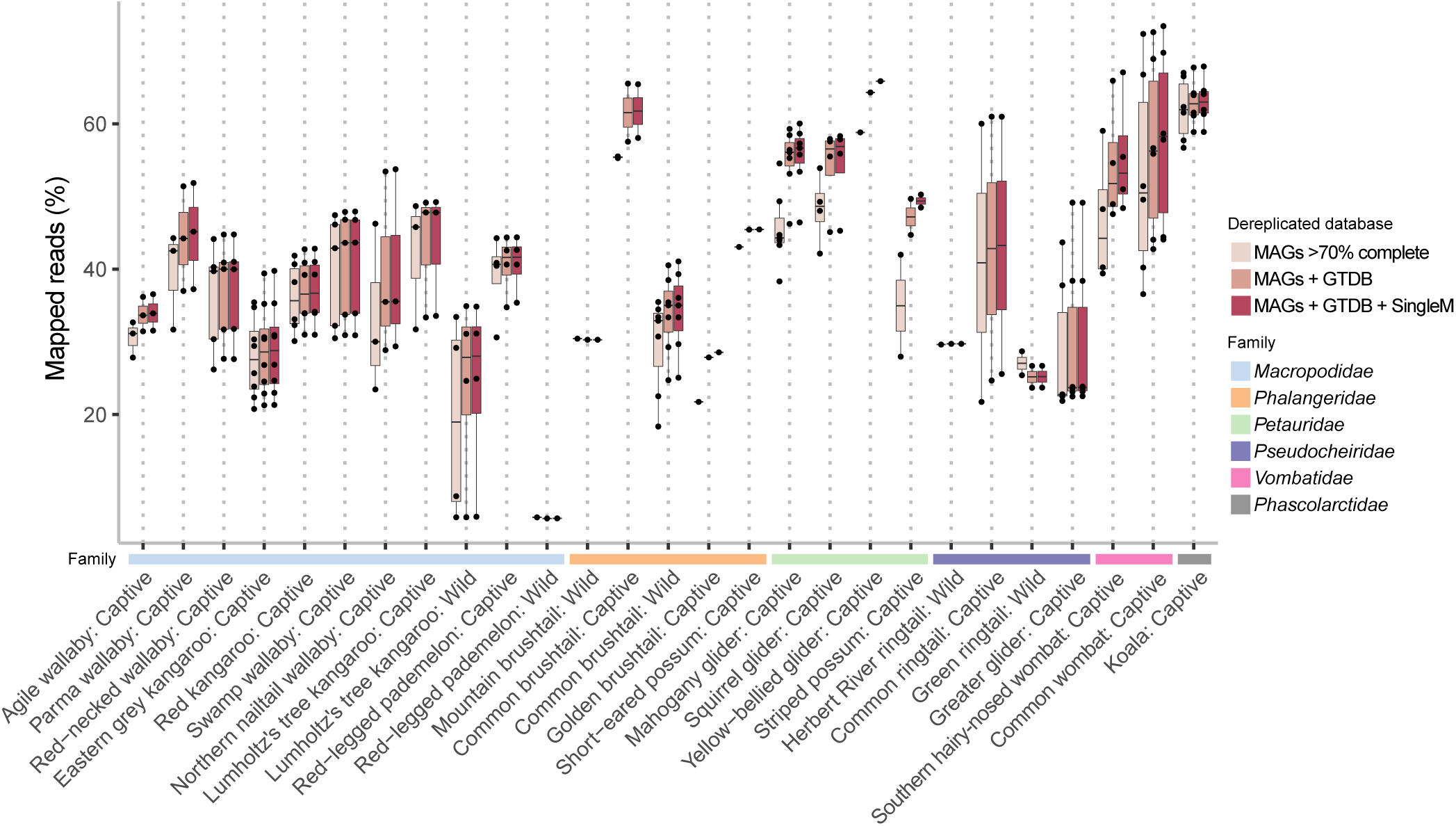
Read recruitment to MAG database incorporating MAGs from the current study ≥70% complete with ≤5% contamination, or study MAGs combined with public genomes selected based on read mapping or marker gene-based profiles. Read mapping proportions based on filtered alignments with ≥95% identity across ≥90% of the read length.

**Fig. S5.**
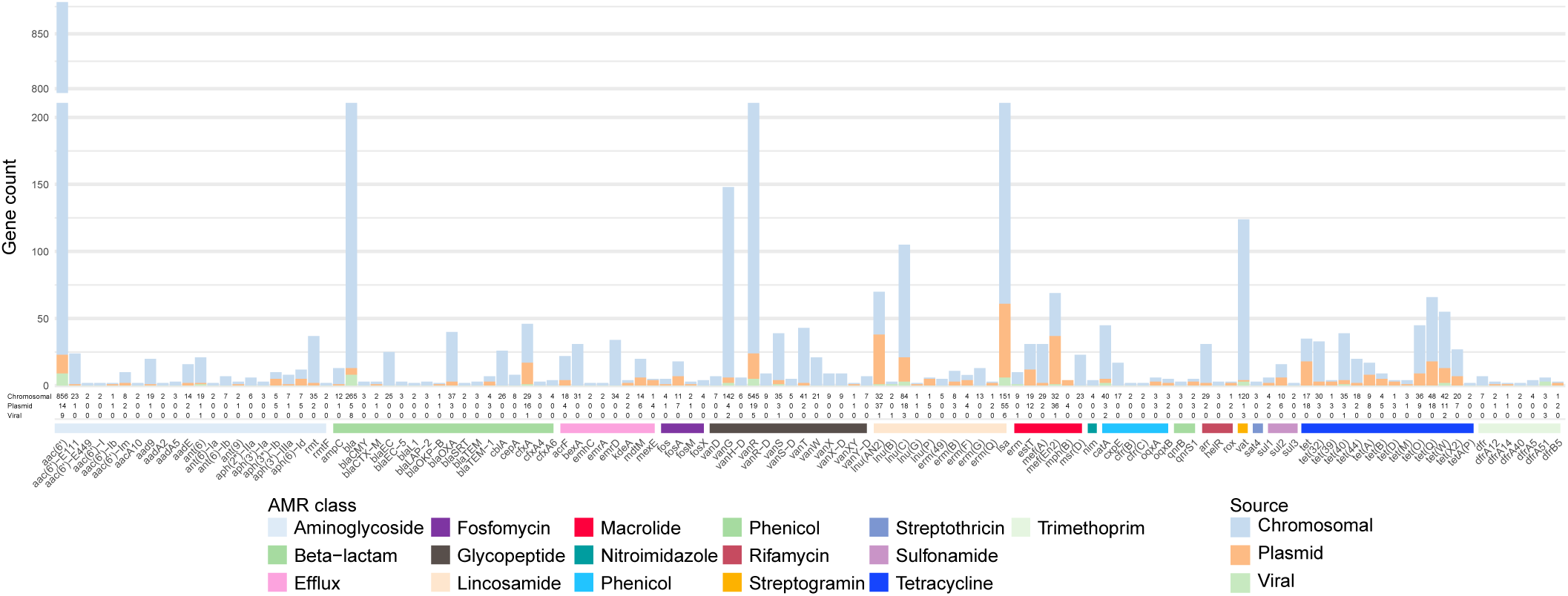
Count of AMR genes identified on all contigs based on AMRFinder search, excluding singletons. Source indicates the Genomad-predicted sequence type of the contig carrying the representative gene sequence.

**Fig. S6.**
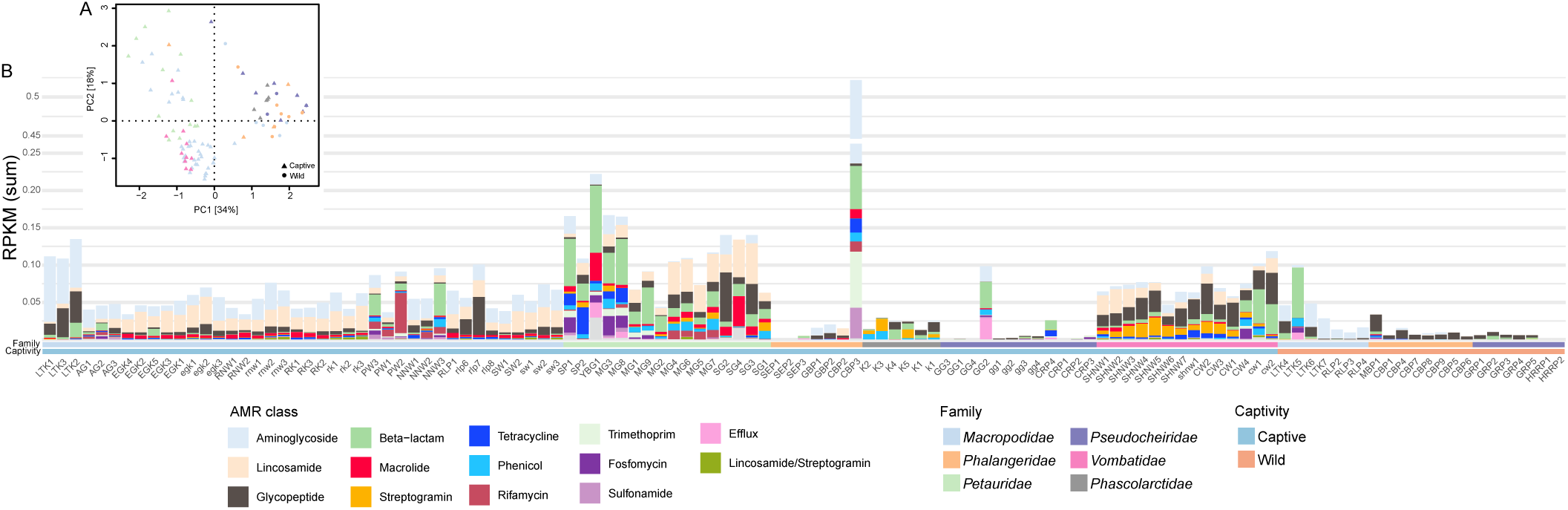
a. PCA based on AMR gene RPKM values. **b** RPKM sum per AMR class per marsupial faecal sample.

**Fig. S7.**
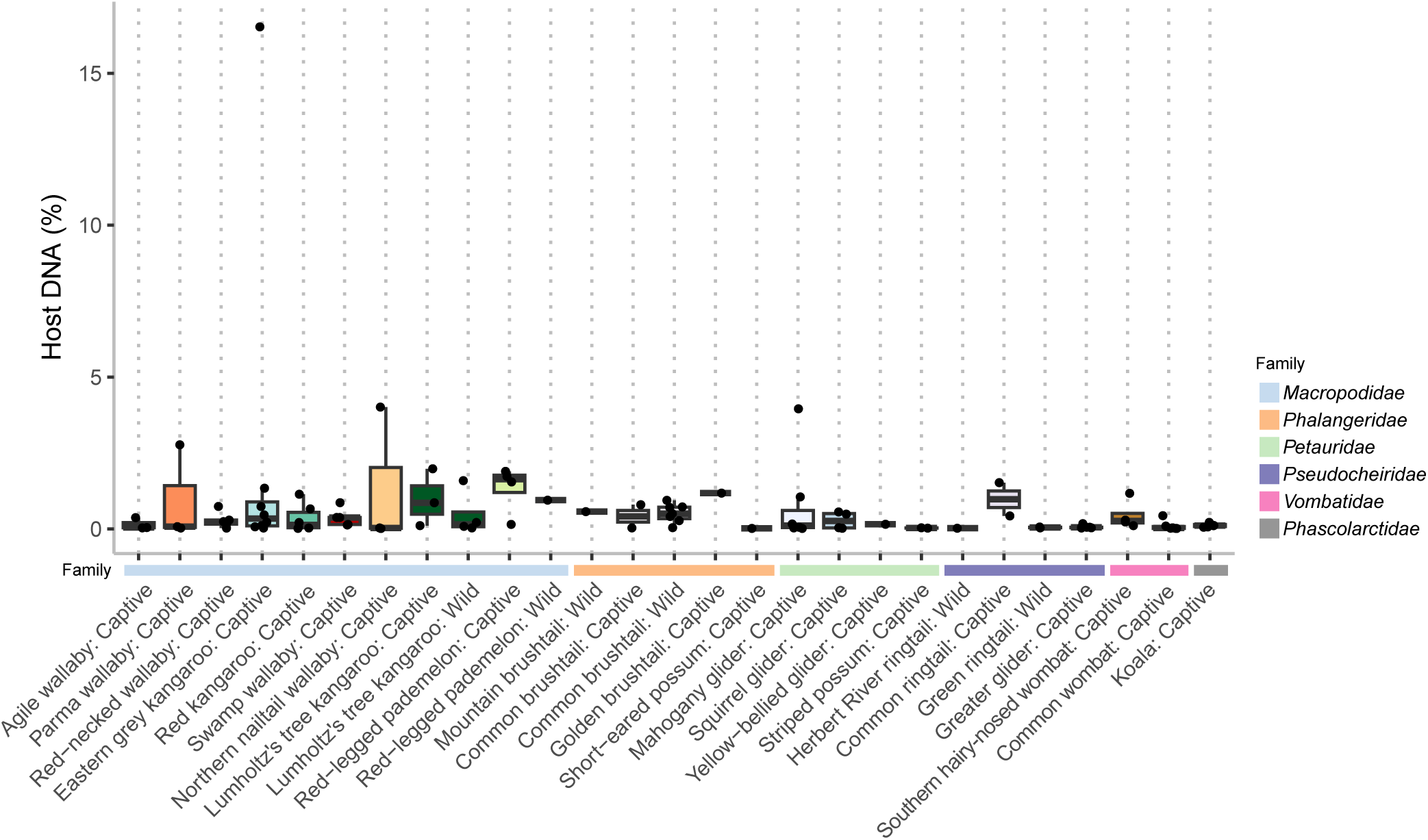
The relative amount of host animal DNA in the faecal metagenomes was estimated by mapping sequencing reads to publicly available marsupial host genomes, including an exon capture dataset obtained from NCBI Genbank (Benson et al., 2013). As found in other animals, host contamination was low (<4% of total reads; Ong et al., 2022), with one conspicuous outlier from an eastern grey kangaroo (∼17%) that may indicate elevated gut epithelial cells in the feces due to inflammation or other dysbiosis (Jiang et al., 2020).

